# Does increasing the dose of relaxation strain augment the sarcomerogenic response to isometric resistance exercise?

**DOI:** 10.64898/2026.10.06.757082

**Authors:** Keenan B. MacDougall, Brian R. MacIntosh, Geoffrey A. Power

**Author notes:** **Corresponding author:**Geoffrey A. Power PhD. Neuromechanical Performance Research Laboratory, Department of Human Health Sciences, College of Biological Sciences, University of Guelph, Guelph, Ontario, N1G 2W1, Canada.

## Abstract

Increases in serial sarcomere number (SSN) have been suggested to reduce muscle strain injury risk and improve muscle mechanical performance, however the environmental loading conditions optimizing sarcomerogenesis are not well known. It has been suggested that the magnitude of lengthening which fibres are subject to upon deactivation (i.e., the “relaxation strain”) may provide a stimulus for sarcomerogenesis. However, no study has attempted to amplify this relaxation-strain stimulus to directly test this possible mechanism. Sprague-Dawley rats underwent six weeks (twice weekly) of isometric plantar flexion training, either sustained near-maximal contractions, or intermittent ‘burst’ type contractions (i.e., ‘contract-relax’) in which each repetition consisted of eight brief high-frequency trains to magnify the overall incidence of relaxation strain occurrences across the training period. Groups were otherwise matched for peak torque and torque-time integral. Following training, the torque-angle relationship adapted similarly between groups (training×group interaction P=0.249), with active torque increasing ∼30-40% across the angles tested (P<0.001). No changes in maximal shortening velocity were observed with training in either group (P=0.577), however peak power output increased by ∼19% in Isometric and ∼49% in Burst (training effect P=0.002; interaction P=0.152). For SSN, the gastrocnemius showed a divergent training response (Lateral gastrocnemius: Burst +∼11%; Isometric +∼2%; interaction P=0.001; Medial gastrocnemius: Burst +∼9%; Isometric +∼3%; interaction P=0.052), while soleus SSN increased similarly between groups (∼2-3%) (training effect P=0.029, interaction P=0.543). Overall, amplifying the relaxation strain stimulus may have a positive effect on stimulating sarcomerogenesis, however further research is needed to clarify the extent of this effect, as well as any underpinning mechanisms.

## 1. INTRODUCTION

In response to resistance training, muscle fibres may increase both their cross-sectional area, as well as in their length, the latter of which is typically due to an increase in the number of sarcomeres in series ^1,2^. Importantly, this process, known as sarcomerogenesis, can have numerous potential benefits. For example, increases in serial sarcomere number (SSN) improves muscle force production at longer muscle lengths ^2,3^, as well as during high speed movements ^4^, resulting in reduced muscle strain injury risk ^5,6^ as well as improved power and sprint performance ^7–10^.

Despite the practical relevance of increasing a muscle’s SSN, the signaling mechanisms that underpin sarcomerogenesis, and the mechanical stimuli that optimize it, remain poorly understood. A recent review ^11^ highlighted this knowledge gap, identifying several candidate mechanical stimuli that may contribute to the mechano-transduction signal underpinning sarcomerogenesis. For example, along with emphasizing the importance of high muscle tension during training ^3^, the potential importance of muscle length in regulating SSN was also highlighted in a study by Uçar et al, ^12^, which reported a 32% increase in SSN in the rat extensor digitorum longus (EDL) in response to isometric training at long muscle lengths, compared to a 12% increase when trained at moderate lengths. Comparatively, Rilling et al., ^13^ recently showed a ∼9% increase in the rat soleus SSN following isometric training at moderate muscle lengths. Interestingly, while Blazevich et al., ^11^ suggested that active lengthening (e.g., eccentric) contractions did not seem to positively impact SSN *per se* (although cf. Refs ^14,15^), they proposed that relaxation strain – defined as the lengthening of muscle fibres back toward resting length as force declines during deactivation ^16^ – may act as a trigger for sarcomerogenesis. Evidence supporting relaxation strain as a potential sarcomerogenic stimulus comes primarily from Butterfield and Herzog ^3^, who found that increases in SSN following six weeks of eccentric training in rabbit tibialis anterior were most strongly associated with peak torque and the fibre relaxation strain. In humans, following ten weeks of eccentric knee extension exercise at varying lengthening velocities, Sharifnezhad et al., ^17^ found that vastus lateralis fascicle length was only significantly increased in the group that trained at the fastest lengthening velocities, a result which they attributed to a period of apparent muscle deactivation occurring at the end of the contractions, thus likely increasing the magnitude of the relaxation strain experienced by the fibres.

Thus, despite being speculated to act as a trigger for sarcomerogenesis, there is a paucity of evidence for the impact that relaxation strain might have in the training response. Furthermore, no study has attempted to amplify the mechanical signal to more directly explore whether relaxation strain imparts a distinct sarcomerogenic stimulus. Therefore, the purpose of this study was to investigate whether a training program which specifically increases the overall number of relaxation strain occurrences across a training period would lead to a stronger sarcomerogenic response relative to a more conventional sustained isometric training program. Based on these previous observations ^3,17^, it was hypothesized that increasing the “training dose” of this relaxation strain signal would lead to greater increases in SSN and associated mechanical performance compared to a conventional isometric training program.

## 2. MATERIALS AND METHODS

### 2.1. Animals

Twenty male Sprague-Dawley rats (Charles River Laboratories, QC, Canada) were used for this study. Rats began the study at ∼12 weeks of age, weighing ∼350–475g. Rats were housed in an environmentally controlled room at 23°C with a 12hr reverse light-dark cycle, and given standard rat chow (Envigo Teklad 2918, Indianapolis, USA) and water *ad libitum*. All procedures were approved by the University of Guelph Animal Care Committee (AUP#4905).

### 2.2. Resistance training protocols

Rats were assigned randomly to one of two training groups (n=10 each). Prior to all training and testing sessions, rats were anesthetized with isoflurane and positioned supine on a heated platform (37°C). Their left leg was shaved and then secured via surgical tape to a foot plate connected to a torque transducer (Model 1305B, Aurora Scientific, Aurora, Canada), with the leg clamped at a knee angle of ∼150° (full extension 180°). Transcutaneous electrical stimulation was applied to the hindlimb of the left leg via a bi-phase electrical stimulator (Model 701C, Aurora Scientific) and a custom-made electrode holder using galvanized steel pins placed over the popliteal fossa and Achilles tendon ^18^. At the beginning of each training or testing session, maximal stimulation current was found using a series of tetanic stimulations (stimulation frequency 100Hz, pulse width 0.1ms, train duration 250ms) at an ankle angle of 70° (full plantarflexion 180°). Stimulation current was increased in increments of 10mA until torque reached a maximum, and this current was then used for the duration of the session.

To isolate the impact of increasing the “dose” of relaxation strain on muscular adaptations, we minimized the potential impact of other training variables. Specifically, peak torque, and torque-time integral ^3,19^ were matched as closely as possible between training groups (Isometric and Burst). During pilot testing, we evaluated various stimulation parameters: pulse frequency, train frequency, pulses per train, and train duration. It was found that the response to a series of brief stimulations (8 trains of 12 pulses at 120Hz, every 0.25s) well-matched the peak torque and torque-time integral of a single train of 80Hz stimulation for 1s (where each response was referred to as a single repetition). Each training set was comprised of 6 repetitions at 5s intervals, and 3min was allowed between sets. For both groups, training was conducted 2× weekly for 6 weeks (12 total training sessions), with 72–96 hours between sessions. Weekly training load was manipulated across the training period, beginning with 3 sets for the first week, and adding 1 set each week. In the 6^th^ week, there were 8 sets of 6 repetitions, but this was reduced to 3 sets for the final session. Figure 1A and 1B shows representative torque-time tracings for a single repetition for the Burst and Isometric conditions, respectively.

**Figure 1.**
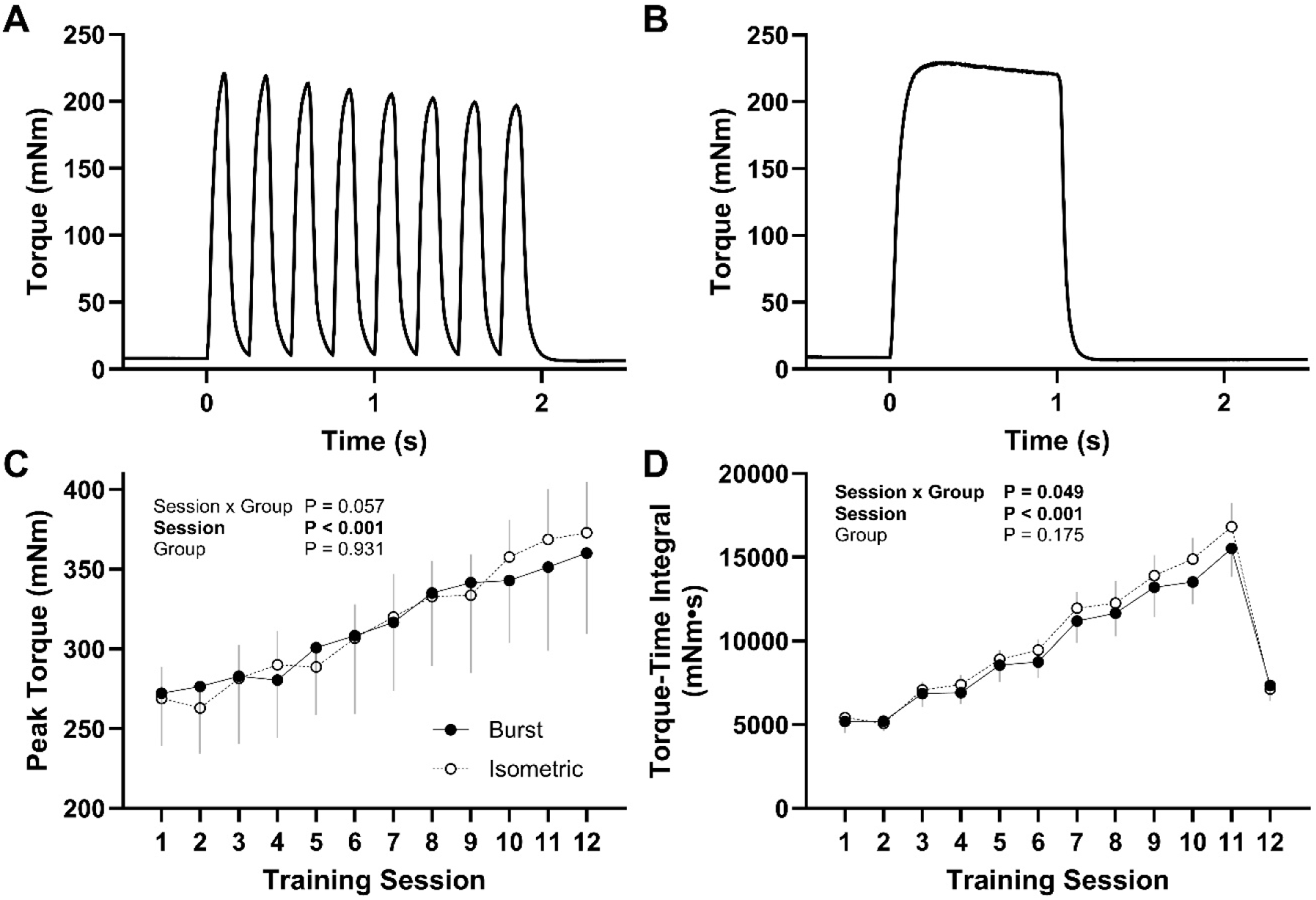
Representative torque traces and training load data for Burst and Isometric training protocols. **A**: representative torque-time trace for a single repetition in the Burst group. **B**: representative torque-time trace for a single repetition in the Isometric group. **C**: Changes in peak torque output measured during the training sessions across the training period. **D**: Changes in torque-time integral per session across the training period. Data is shown as mean ± SD. Burst n=10, Isometric n=9.

### 2.3. Mechanical testing

All mechanical testing was performed 48–72 hours before the commencement of the training program, and 48–72 hours following the training program. Active torque-angle relationships were constructed with isometric contractions using 100Hz stimulation for 500ms, at ankle angles of 70°, 75°, 80°, 90°, 100°, and 110°. Contractions were performed in a randomized order, with 2min of recovery between. For each contraction, active torque was calculated as the peak torque minus the passive torque occurring at the onset of stimulation. While this method underestimates the active torque produced when passive force is high ^20^, it should have minimal impact across the angles tested here.

Torque–angular velocity relationships were constructed using a series of afterloaded isotonic contractions. The protocol began with an isometric contraction at an ankle angle of 70° (the starting position for the isotonic contractions), using a stimulation frequency of 200Hz for 500ms. Subsequently, 200ms isotonic contractions were performed in a randomized order against loads corresponding to 5%, 15%, 25%, 35%, 45%, 55%, 65%, 75%, and 85% of the peak torque generated during the isometric reference contraction, with 2 min of recovery between contractions. To allow a more direct comparison in angular velocity improvements at a given load, during post-testing, it was decided to construct the torque-angular velocity relationships using the same absolute isotonic loads as that used during pre-testing, rather than to use the same relative loads. As such, to ensure sufficient resolution in the high-load region of the torque-angular velocity relationship for curve fitting, while accounting for training-induced improvements in maximal torque output, three additional contractions were performed at 95%, 105%, and 115% of the pre-training peak isometric torque. On average, the 115% load corresponded to 84±5 % of the post-training peak isometric torque value. All torque and length data were collected at a sampling frequency of 2000 Hz, using commercial software (DMC version 5.5, Aurora Scientific) and subsequently exported to a personal computer for analysis.

The angular velocity for each afterloaded contraction was measured as the highest value of a 20ms moving average of the derivative of the foot plate position signal, with the corresponding torque value taken as that over the same time interval. Power output for each contraction was then calculated as the product of torque and angular velocity. For curve fitting, torque and angular velocity data from the dynamic contractions were fit to the Hill equation:

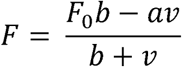

Where *F* is the torque produced, *v* is the angular velocity, *F_0_* is the predicted maximal isometric torque, and *a* and *b* are constants ^21^. From these parameters, the predicted maximal shortening velocity *V_max_* as well as the curvature constant *a/F_0_* was calculated. Curve fitting was done in R version 4.5.2 using nonlinear least squares regression with the Levenberg–Marquardt algorithm. The average *r*^2^ value for the curve fits across pre– and post-testing was 0.994±0.005.

### 2.4. Morphological analyses

24-48 hours following post-training mechanical testing, rats were euthanized using isoflurane and CO_2_ inhalation. Both left and right hindlimbs were skinned, amputated, and fixed the ankle and knee at 90° in 10% phosphate buffered formalin for 4 weeks. Following fixation, the soleus, medial gastrocnemius, and lateral gastrocnemius of each limb were dissected, blotted dry, and weighed. The muscles were then rinsed with phosphate-buffered saline and digested in 30% nitric acid for ∼6 hours prior to being stored in glycerol. For each muscle, 6 fascicles were teased out (2 from the proximal region, 2 from the middle region, and 2 from the distal region) for serial sarcomere number determination. Each fascicle was placed on a glass microslide, and sarcomere lengths were measured at six locations across the length of the fascicle using laser diffraction with a 5-mW diode laser (∼1 mm beam diameter, 635 nm wavelength) (Coherent, Santa Clara, CA, USA) and custom LabVIEW program (National Instruments, Austin, TX, USA). Photographs of the fascicles were then taken using a digital camera mounted on a tripod, and from these, fascicle lengths were measured using ImageJ software (version 1.54), with measurements calibrated to a ruler in plane with the fascicles. For each fascicle, serial sarcomere number was calculated as the measured fascicle length divided by the average of the 6 measured sarcomere lengths. Finally, overall average sarcomere length, average fascicle length, and average SSN was then calculated from the 6 fascicles to obtain a single representative value for each muscle. From these averaged values, the physiological cross-sectional area (PCSA) for each muscle was calculated as:

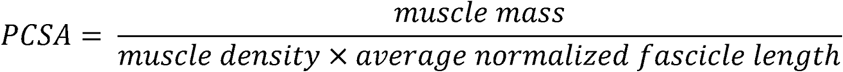

Where muscle mass was taken as the measured wet weight of the muscle, muscle density was assumed to be 1.112 g/cm^3^ ^22^, and normalized fascicle length for each fascicle was calculated as:

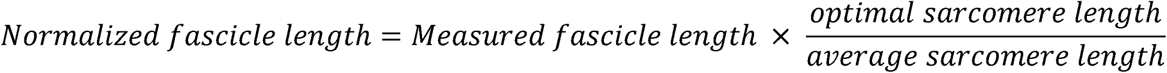

Where optimal sarcomere length was assumed to be 2.4μm ^23^.

### 2.5. Statistical analyses

Data are presented as mean ± standard deviation. Statistical analyses were conducted using Graphpad Prism version 11.1.0 (Graphpad Software, Boston, USA). To confirm that training load was properly matched between groups, the peak torque output, as well as the total torque-time integral across the training sessions was compared using a two-way repeated measures Analysis of Variance (ANOVA) (Training Session [1-12] × Group [Burst, Isometric]). Alterations in the active torque-angle relationships were analyzed using a three-way repeated measures ANOVA (Training [pre, post] × Angle [70°-110°] × Group [Burst, Isometric]). Additionally, a two-way repeated measures ANOVA was done on the percentage increase in active torque across the angles tested (Angle [70° – 110°] × Group [Burst, Isometric]). Training induced changes in the measured peak power output, velocity at peak power, *V_max_*, and *a/F_0_* were compared using a two-way repeated measures ANOVA (Training [pre, post] × Group [Burst, Isometric]). Finally, training induced changes in morphological variables (average SSN, fascicle length (FL), sarcomere length (SL), wet weight, and PCSA) were compared using a two-way repeated measures ANOVA (Training [untrained leg, trained leg] × Group [Burst, Isometric]). Post hoc comparisons were made using a Bonferroni correction applied. Normality of the data was confirmed using a Shapiro-Wilk test, and a Greenhouse-Geisser correction was applied where appropriate. For select variables of interest, within group effect sizes (ES) were reported as Hedge’s *g* for paired differences, calculated as the mean of the post-training – pre-training (or trained – untrained) difference scores divided by the standard deviation of the difference scores, with a correction factor applied for small sample sizes ^24^. ES were interpreted as small effect: ES>0.2; medium effect: ES>0.5; large effect: ES>0.8 ^25^. During the training period, one rat in the Isometric group died, so the final sample size for all analyses was n=10 for Burst and n=9 for Isometric. Significance was set at P<0.05.

## 3. RESULTS

### 3.1. Training data

Across the 6-week training period, peak torque amplitude during the training sessions increased similarly in both groups (session effect P<0.001), with no group effect (P=0.931) or session×group interaction (P=0.057) (Figure 1C). Collapsed across both groups, peak torque increased by 35±11% on average from the first to last training session. The torque-time integral, used as a measure of training volume, expectedly changed across sessions as a result of manipulating the weekly set volume (session effect P<0.001). While there was a significant session×group interaction (P=0.049), post hoc comparisons did not reveal any differences between groups within any training session (P≥0.436) (Figure 1D). Across the entire training period, there was no difference in the overall torque-time integral between Burst (113876±11621mNm·s) and Iso (120290±7379mNm·s) (P=0.175).

### 3.2. Morphological adaptations

For the soleus, SSN was greater in the trained versus untrained leg for both Burst (+1.8±2.7%, ES=0.59) and Iso (+3.0±5.6%, ES=0.48) (P=0.029), with no differences between groups (P=0.716), or training × group interaction (P=0.543) (Figure 2A). Soleus wet weight and PCSA were smaller in the trained leg versus the untrained leg (both P<0.001), with no differences between groups (P=0.528) or interaction effects (P=0.253) (Figure 2D and 2G. Relative to the untrained leg, wet weight was 7.6±11.8% lower in the trained leg in Burst (ES=-0.67), compared to 14.5±8.0% smaller in Iso (ES=-1.58), with similar relative differences observed for PCSA. Soleus FL and SL measurements from which SSN values were calculated from are shown in Supplemental Figure 1A and 1D.

**Figure 2.**
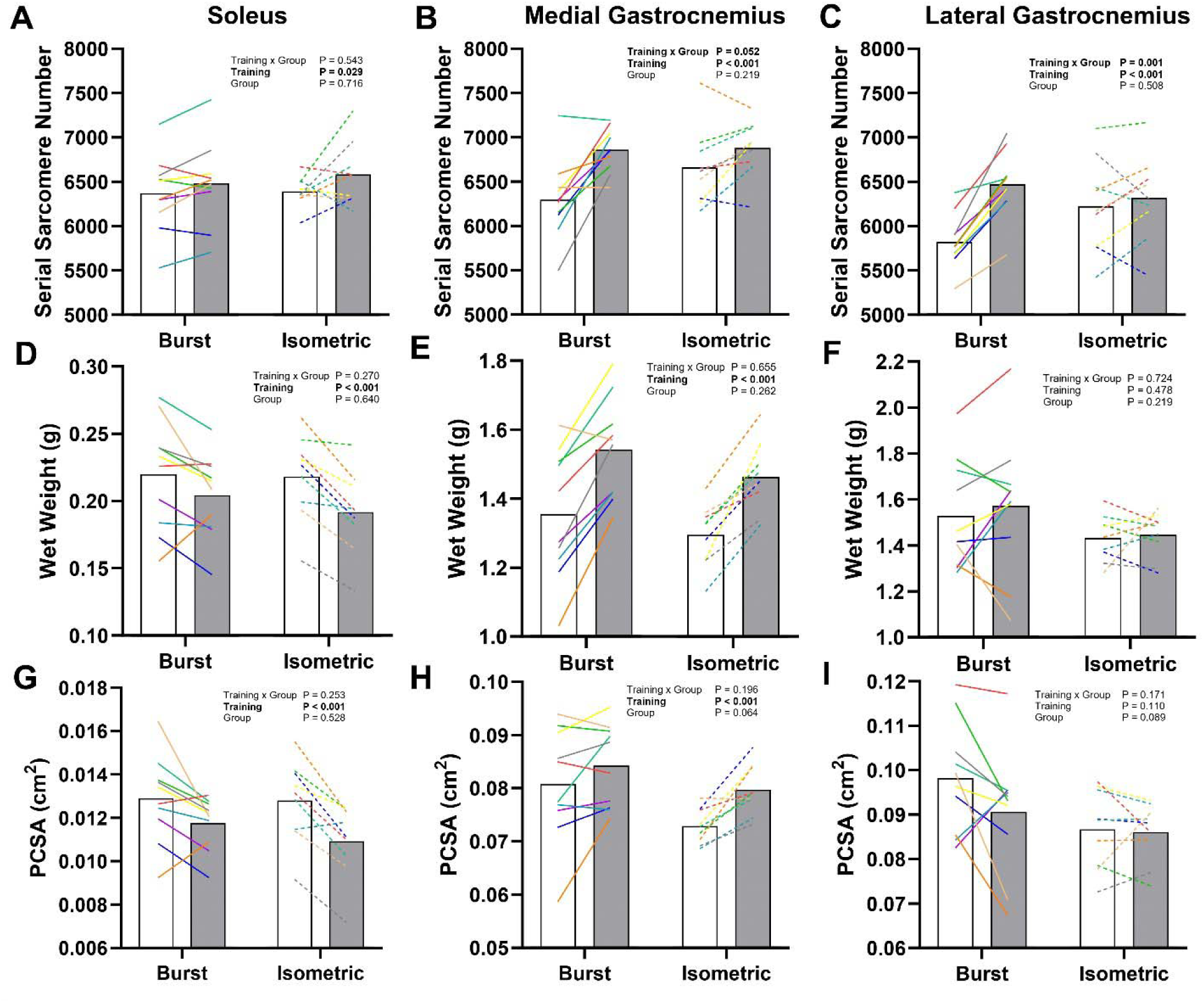
Differences in serial sarcomere number, wet weight, and physiological cross-sectional area (PCSA) between the untrained leg (white) and the trained leg (grey) for soleus (Panels **A**, **D** and **G**), medial gastrocnemius (Panels **B**, **E** and **H**), and lateral gastrocnemius (Panels **C**, **F**, and **I**). Data is shown as mean values, with individual data represented by the coloured lines (solid = Burst, dashed = Isometric). Burst n = 10, Isometric n = 9. Note that the training×group interaction for MG SSN was trending towards significance (P = 0.052).

SSN for the MG was greater in the trained leg relative to the untrained leg (P < 0.001), with no group effect (P=0.219), but a trend for a training × group interaction, (P=0.052) (Figure 2B). SSN was 9.3±7.0% greater in the trained leg (ES=1.27) in Burst, compared to 3.5±4.6% in Iso (ES = 0.66). The wet weight of the trained leg was greater than the untrained leg (P<0.001), with no effect of group (P=0.262) or interaction (P=0.655) (Burst: +14.7±9.0%, ES=1.65; Iso: +13.1±6.5%, ES=1.95) (Figure 2E). MG PCSA revealed an effect of training (P<0.001), with no differences between groups (P=0.067) or interaction effects (P=0.196) (Burst: +5.2±9.4%, ES=0.52; Iso: +9.5±6.1%, ES = 1.39) (Figure 2H). MG FL and SL measurements are shown in Supplemental Figure 1B and 1E.

In the LG, there was a training×group interaction for SSN (P=0.001), whereby post hoc comparisons revealed greater SSN in the trained versus untrained leg in Burst (+11.2±4.5%, ES = 2.23), whereas the difference in Iso (+1.7±5.8%, ES = 0.24) was not significant (P > 0.999). In comparing across groups, there were no differences in SSN in both the untrained (P=0.196) or the trained leg (P>0.999) (Figure 2C). Finally, no significant training, group, or interaction effects were observed for either wet weight (P≥0.219), or PCSA (P≥0.089) of the LG (Figure 2F and 2I, respectively). LG FL and SL measurements are shown in Supplemental Figure 1C and 1F.

### 3.3. Mechanical performance changes

When exploring alterations in the active torque-angle relationship, a three-way repeated measures ANOVA revealed an angle×training interaction (P<0.001), where torque increased across all angles tested (P<0.001), with the absolute increase being greater at longer lengths (Figure 3). Averaged across all angles tested, the Burst group improved active torque production by 39.8±11.7% (ES=3.1), while the Isometric group improved by 36.7±8.8% (ES=4.5).

**Figure 3.**
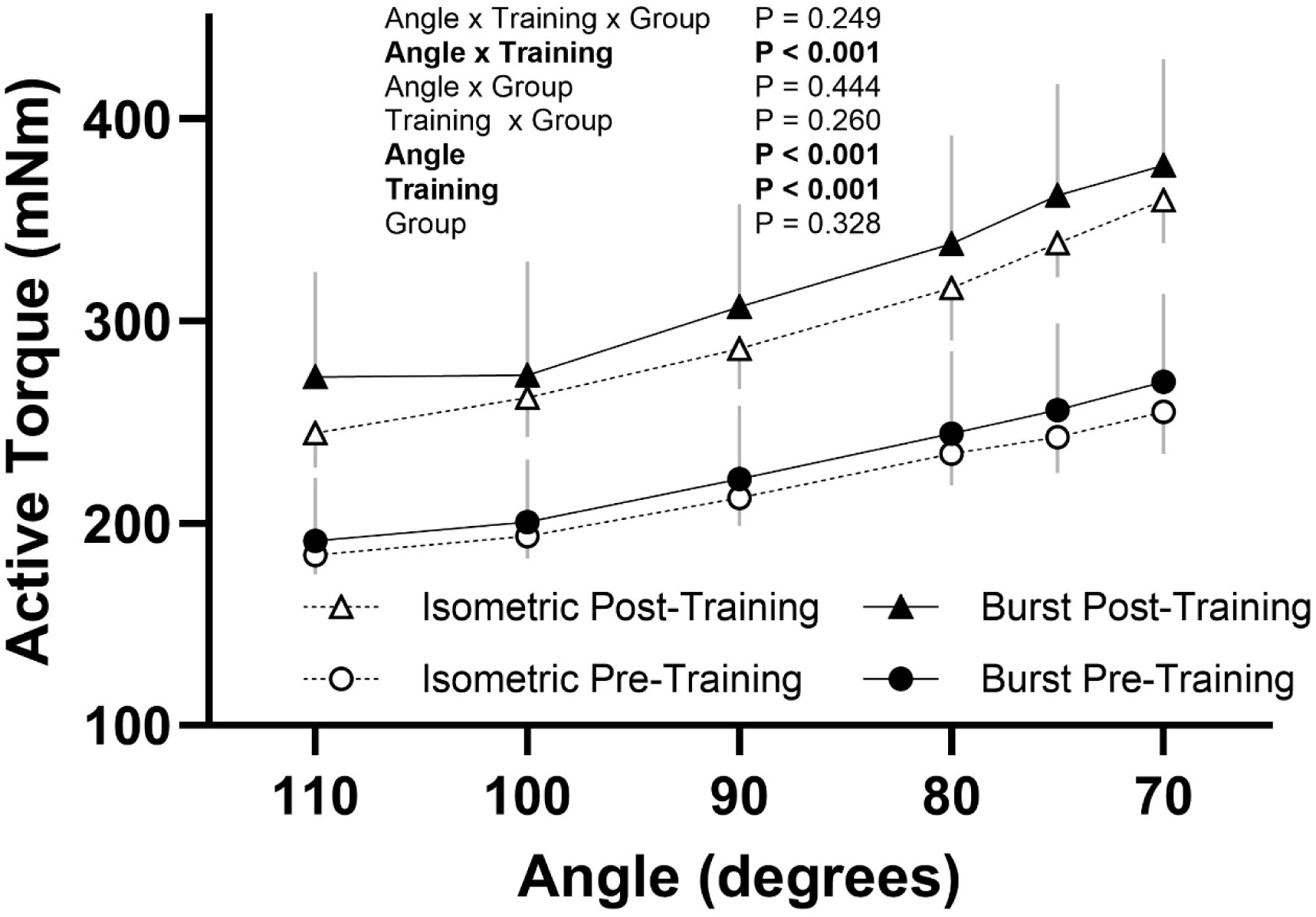
Active torque output across the ankle angles tested pre-training (circles) and post-training (triangles) in the Burst group (filled) and Isometric group (open). Data are shown as mean ± SD. Burst n = 10, Isometric n = 9.

Alterations in the torque–angular velocity and power–angular velocity relationships are shown in Figure 4A and 4B, respectively. Following training, peak power was increased in the Burst group (+48.5±43.4%, ES=1.02) as well as in the Isometric group (+18.7±29.1%, ES=0.51) (P<0.001), with no statistically significant differences between groups (P=0.344) or training × group interaction (P=0.152) (Figure 4C). There was no effect of training (P=0.577), group (P=0.618), or training×group interaction (P=0.442) for *V_max_* (Figure 4D). No training, group, or interaction effects were observed for velocity at peak power (P≥0.515) (Burst pre-training: 648±116deg/s; Burst post-training 621±107deg/s; Isometric pre-training 616±155deg/s; Isometric post-training 607±101deg/s), nor for the curvature parameter *a/F_0_*(P≥0.055) (Burst pre-training: 0.76±0.31; Burst post-training 0.61±0.33; Isometric pre-training 0.55±0.16; Isometric post-training 0.47±0.14).

**Figure 4.**
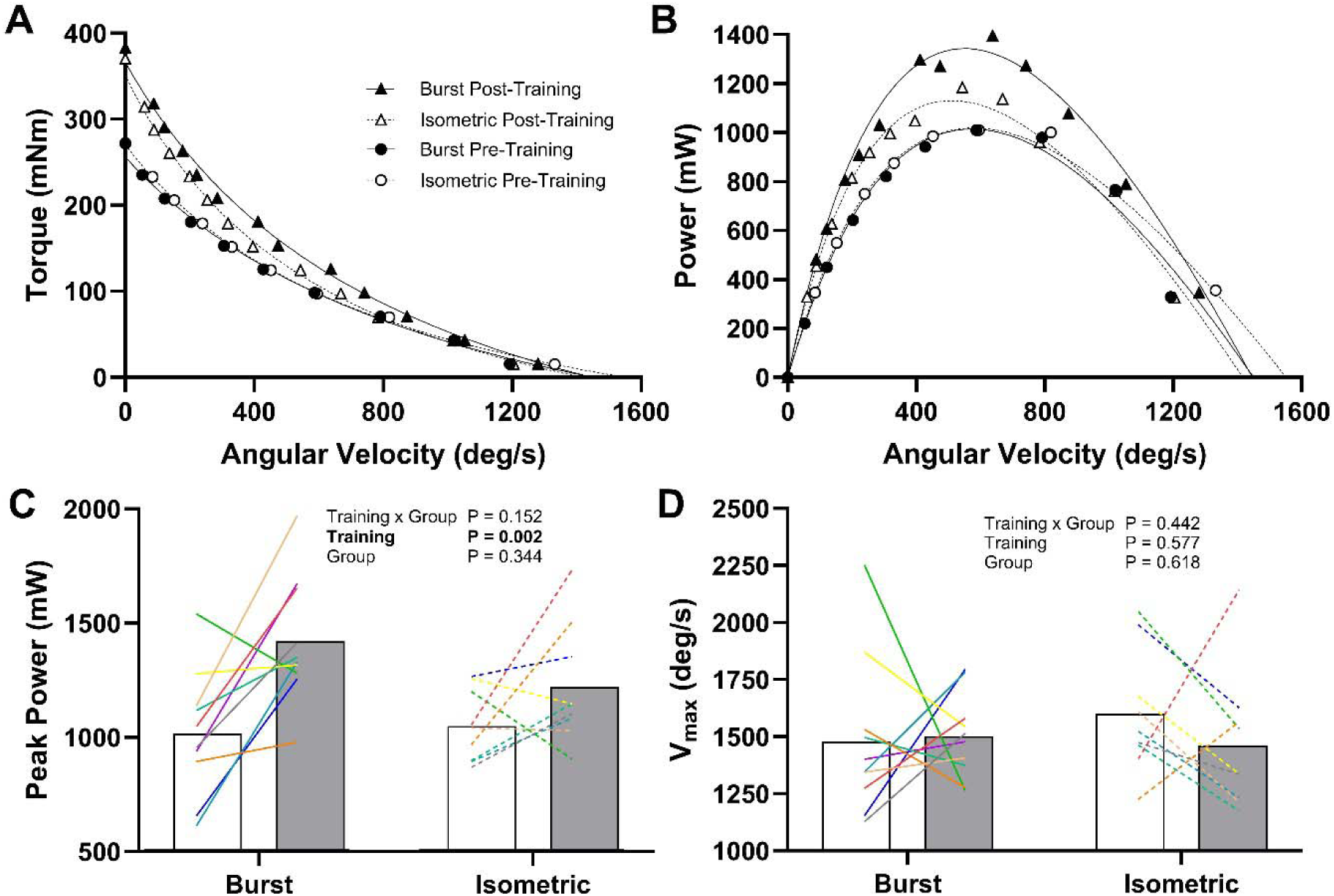
Alterations in the torque–angular velocity–power relationship following training. Panel **A**: Torque – angular velocity data tested pre-training (circles) and post-training (triangles) in the Burst group (filled) and Isometric group (open). Panel **B**: Power – angular velocity relationship. Fitted curves shown were calculated from the mean values for the Hill parameters. For Panels A and B, data points are shown as mean values, with error bars removed for clarity. Panel **C**: Alterations in peak power output from pre-training (white) to post-training (grey). Panel **D**: Alterations in *V_max_*. For Panels C and D, data is shown as mean values, with individual data represented by the coloured lines (solid = Burst, dashed = Isometric). Burst n=10, Isometric n=9.

## 4. DISCUSSION

The present study examined the impact of increasing the “training dose” of the relaxation strain stimulus on training-induced increases in serial sarcomere number, as well as improvements in mechanical performance. In both gastrocnemius muscles, there was evidence for relatively greater longitudinal growth following a training program in which the relaxation strain stimulus was amplified (i.e., intermittent isometric) compared to a more conventional sustained isometric training program. Alternatively, we did not detect any statistically significant group differences in the training responses for the soleus, nor for any of the mechanical variables we explored (although it may be noted that the effect size for peak power was twice as large in the Burst compared to the Isometric group). Therefore, it is possible that the mechanical signal created as muscle fibres lengthen during relaxation may play an important role in the sarcomerogenic response to resistance training. More research is needed to corroborate these findings.

### 4.1. Morphological adaptations with training

In the present study, the soleus adapted to training similarly between groups, increasing SSN by ∼2-3% on average. The gastrocnemius muscles, on the other hand, showed more divergent responses, such that for the lateral gastrocnemius, the increase in SSN was significantly larger in the Burst group (∼11%) compared to the Isometric group (∼2%), while in the medial gastrocnemius, the difference between groups (∼9% versus ∼3% for Burst and Isometric, respectively) was nearing the threshold of significance (P = 0.052), and effect sizes were notably larger in Burst for both muscles (Figure 2). The relatively greater training response in two of the three muscles tested would seem to warrant some consideration that the mechanical stimulus arising from the relaxation strain may exert a positive impact on the regulation of SSN. In comparison to these values, Rilling et al.,^13^ reported a 9% increase in soleus SSN following an isometric training protocol (differing in volume and frequency from the Isometric protocol employed here), however they reported no increases in MG SSN. Uçar et al., ^12^ compared the response to an isometric training protocol at long versus moderate muscle lengths in the rat EDL, and found a 32% increase in SSN when trained at long lengths, versus a 12% increase when trained at moderate lengths. However, the post-training SSN did not differ between groups. Across various other training studies in the rat plantarflexors, increases in soleus SSN have ranged from ∼2% up to ∼17%, while the MG has ranged from no change to up to an ∼8% increase ^26–29^. The only other study which included the lateral gastrocnemius reported no significant changes in SSN following a maximal eccentric training program ^28^. Changes in wet weight following resistance training have been similarly inconsistent in both soleus and MG, ranging from no change up to ∼13-14% ^26–29^. Curiously, while in the present study we saw a similar ∼14% increase in MG wet weight in the trained leg across both groups, we observed an apparent reduction in soleus wet weight with training. While also considering the relatively small improvements we observed in soleus SSN, the reason for this apparently maladaptive response in terms of overall muscle mass is not entirely clear. While speculative, one potential explanation is that the session training volume in the current study reached substantially higher volumes (8 sets) compared to previous studies (3 sets) ^26–29^. This added training load may have resulted in persistent fatigue or muscle soreness of the trained limb, causing the rats to put relatively more load onto the untrained limb during the non-training periods. Considering the soleus is a postural muscle that is highly active even during quiet standing (unlike the gastrocnemius) ^30^, it is possible that off-loading the trained side during quiet periods may have paradoxically shifted the training response to be relatively more in favor of the untrained soleus. Additionally, because MTU or fascicle length was not measured or controlled during training, reliance on a fixed ankle joint angle may have resulted in differences in training muscle lengths due to variation in rat leg size across studies. Future work should directly measure and control for muscle length during training.

Throughout the literature utilizing animal resistance training models, there appears to be substantial variability in the morphological training responses observed (e.g., SSN, wet weight, PCSA), both across different muscles for a given training program, as well as across training programs for a given muscle. On the whole, however, our results are in general alignment with previous studies ^26–29,31^. This heterogeneity in training response is perhaps not surprising, given the many training-related factors (e.g., intensity, volume, frequency, joint vs. muscle excursion) ^3,19,32–34^ as well as physiological factors (e.g., animal strain, sex, age) ^26–28,35,36^ that may ultimately impact the morphological response to training, and may obfuscate any interpretations made when comparing across studies. In the current study, these factors were matched as closely as possible between groups, and so the fact that we observed some (limited) evidence of an improved response to training in the Burst group would suggest that it was the stimulus arising from the added relaxation strain, and not one of these other potential confounding factors (e.g., torque differences), that underpinned this effect.

### 4.2. Alterations in the torque-angle relationship with training

One of the characteristic functional implications of an increase in SSN is a rightward shift in the active force-length or torque-angle relationship. For example, Butterfield and Herzog ^3^ compared three separate eccentric exercise protocols in the rabbit tibialis anterior, and found that the two protocols which resulted in more robust increases in SSN showed clear rightward shifts in optimum joint angle for torque production, and this shift occurred concomitantly with a length-dependent increase in the percentage change in torque output following training (i.e., torque output improved to a relatively greater extent at longer versus shorter muscle lengths). Alternatively, Butterfield and Herzog ^3^ observed that the optimum angle was unchanged following the protocol which showed only marginal changes in SSN, and no clear length dependence of the percentage change in torque following training was apparent. In the present study, the ankle and testing apparatus did not permit sufficient range of motion to stretch the muscle onto the descending limb of the torque–angle relationship, and therefore prevented us from observing any shifts in optimum angle (Figure 3). While the angle×training interaction we observed indicated that the *absolute* improvements were greater at the longer muscle lengths, this was not borne out when expressing the improvement as a percentage increase, as done by Butterfield and Herzog ^3^ (Supplementary File Figure 2). Nevertheless, considering that the knee was flexed in our experimental setup, it might be that the more substantial increases in SSN we observed in the bi-articular gastrocnemius compared to the mono-articular soleus would not translate to large-scale changes in the joint-level torque-angle relationship. Therefore, (and given the previously mentioned limitations in the range of motion of the experimental setup), these results are perhaps not surprising.

### 4.3. Alterations in the torque-angular velocity-power relationship with training

According to the hyperbolic force-velocity relationship, increases in peak power may come about due to increases in maximal isometric force, increases in the curvature parameter *a/F_0_*reflecting decreased curvature, and/or increases in maximal shortening velocity. Radial growth (e.g., an increase in PCSA) would theoretically function to improve peak power output due to its positive impact on maximal isometric force, without altering the velocity at peak power. On the other hand, an increase in SSN would theoretically function to improve peak power via its positive effects on maximal shortening velocity, concomitant with a rightward shift in the velocity at peak power. In the present study, we observed significant improvements in peak power output, nearing 50% in the Burst group and 20% in the Isometric group, although statistically these were not different (Figure 4). Contrary to our hypothesis, however, we observed no change in either *V_max_* or velocity at peak power (although it might be noted that the Burst group had proportionally more “positive responders” in *V_max_*(6 of 10 versus 2 of 9 for Isometric; Figure 4)). Combined with the more robust increase in peak power output in the Burst group, these data may be taken as evidence for the potential superiority of this training protocol versus conventional isometric training. However, there are clearly more factors at play in determining the improvements in dynamic muscle performance than we were able to measure here.

Interestingly, despite commonly reporting significant improvements in SSN and isometric torque output following training, many of the previously mentioned studies employing various resistance training programs in the rat hindlimb ^13,27,28^ have failed to detect improvements in V_max_, and peak power output. Thus, there may be other adaptations occurring during the training period that are serving to counteract the theoretically positive changes in *V_max_* and peak power output that may be expected with an increase in SSN. As one consideration, there is ample evidence that resistance training generally leads to a faster-to-slower fibre subtype conversion in both humans (i.e., IIx to IIa) ^37^, and in rats (i.e., IIb to IIx) ^38^. Notably, maximal shortening velocities in the intermediate rat IIx fibres have been reported to be in the range of ∼20% lower ^39^ to ∼50% lower ^40^ than that of faster IIb fibres. While it may be argued that theoretically, whole muscle *V_max_* should relate to the *single* fastest motor unit within the muscle ^41^, in practice, it is a value extrapolated from the measured force and velocity data, which would be expected to depend on fibre type distribution. As such, if there were to be significant IIb to IIx shifts following training, this adaptation could conceivably have a large tempering effect on any SSN-induced improvements in the angular velocity at peak power and the apparent *V_max_*.

### 4.4. Putative mechano-transduction mechanisms for sarcomerogenesis

While the mechanical and molecular signaling events underlying skeletal muscle remodeling in response to resistance training is an ongoing area of research ^42–45^, here we will briefly cover several candidate mechano-transduction mechanisms for sarcomerogenesis. Firstly, it may be noted that while muscle damage has historically been theorized to play a key role in sarcomerogenesis ^46,47^ (as well as in underpinning the role of relaxation strain specifically ^17^), recent research suggests that damage is likely not causal in initiating longitudinal remodeling ^48^ (and see Ref. ^28^). Alternatively, Franchi et al., ^49^ showed in humans that activation of focal adhesion kinase (FAK), a protein localized within the costamere, was greater following eccentric versus concentric exercise, and was significantly correlated with the training induced regional increases in fascicle length. Transmembrane proteins linked with focal adhesion complexes are able to sense tension within the sarcolemma ^50^, such as that which would be expected when the muscle is stretched to longer lengths, and thus may be important mediators for sarcomerogenesis. Given the viscoelastic nature of muscle ^51^, whether this FAK-based mechanism is also sensitive to rates of stretch deserves some consideration (e.g., see Burkholder ^52^ as well as Sharifnezhad et al., ^17^). Here, it might be speculated that any impact of relaxation strain may be mediated by its impact on inducing high fibre stretch velocities as the tendon recoils during deactivation. Although not mutually exclusive with a costamere-related mechanism, there is also evidence that the tension experienced by the sarcomeric “spring-like” protein titin plays a primary role in the mechano-sensing process leading to sarcomerogenesis ^53^, highlighted by findings in which artificially increasing titin stiffness via excising a portion of its spring region led to a ∼30% increase in SSN ^54^. Conceivably, the “membrane-biased” FAK signaling mechanism and the “sarcomere-biased” titin signaling mechanism may also interact, considering the potential for lateral force transmission to occur between the Z-lines (of which titin is attached to ^55^) and the costameres ^56^. In relating this titin-based mechanism to the present study, it may be speculated that the “Burst” experimental paradigm is analogous to a series of shortening-stretch cycles, of which, notably, titin tension has been suggested to mediate the relatively increased force observed following a shortening-stretch cycle compared to an isometric contraction at the same final length ^57^ (although cf. Ref ^58^).

## 5. CONCLUSION

Longitudinal muscle growth is a key adaptive response to loading, with important functional benefits for both athletes as well as the general population. The present study was the first to systematically explore the impact of “relaxation strain” on the sarcomerogenic response, which has been suggested to act as a trigger for sarcomerogenesis. Compared to a more conventional isometric resistance training protocol, some of our morphological and performance metrics may point to a positive effect from increasing the “training dose” of this stimulus, however whether this presents a practically meaningful improvement in the training response deserves further investigation. We surmise that if relaxation strain does offer a unique stimulus for sarcomerogenesis, it may be mediated by a transiently increased strain of titin during the period of deactivation, as cross-bridge force declines, and the elastic tendon recoils back to its resting length. Given our findings, however, more research is needed to verify these hypotheses.

## ACKNOWLEDGEMENTS

This work was supported by the Natural Sciences and Engineering Research Council of Canada under Grant RGPIN-2024-03782 and RGPIN-03438-2023.

## AUTHORS’ CONTRIBUTIONS

KBM conceived of and designed the study, performed the experiments, analyzed the data, prepared figures, and drafted the manuscript. GAP conceived of and designed the study and interpreted results of the experiments. BRM interpreted results of the experiments. All authors edited and revised the manuscript. All authors have read and approved the final version of the manuscript and agree with the order of presentation of the authors.

## DATA AVAILABILITY STATEMENT

Data generated or analyzed during this study are available from the corresponding author upon reasonable request.

## FIGURES

**Supplemental Figure 1.**
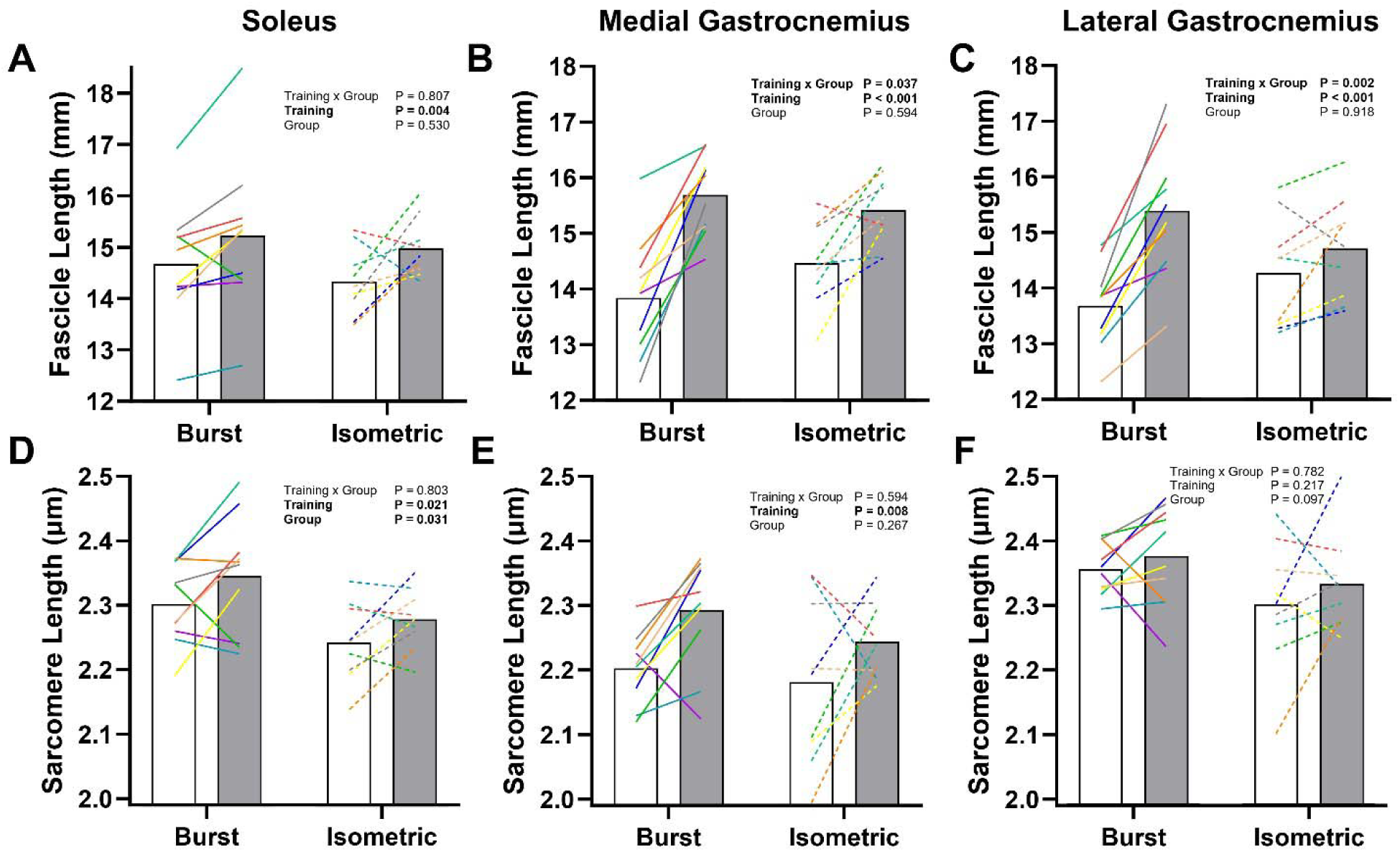
Differences in fascicle length and sarcomere length between the untrained leg (white) and the trained leg (grey) for soleus (Panels **A** and **D**), medial gastrocnemius (Panels **B** and **E**), and lateral gastrocnemius (Panels **C** and **F**). Data is shown as mean values, with individual data represented by the coloured lines (solid = Burst, dashed = Isometric). Burst n = 10, Isometric n = 9. **Results:** In the soleus, there was a significant effect of training on fascicle length (P = 0.004) with no differences between groups (P = 0.530) or any interaction effect (P = 0.807) (Burst: +3.7 ± 4.5%, ES = 0.73; Iso: +4.7 ± 6.2%, ES = 0.66). Soleus sarcomere length was longer in the trained leg versus the untrained leg (P = 0.021), as well as being longer in Burst compared to Iso (P = 0.031), with no interaction effect (P = 0.803). In the MG, a training × group interaction (P = 0.037) revealed differences in the trained leg versus the untrained leg in both Burst (+13.8 ± 7.6%, ES = 1.82; P < 0.001), as well as in Iso (+6.7 ± 5.7%, ES = 0.1.08; P = 0.018). There were no differences in fascicle length between Burst and Iso in the untrained leg (P = 0.420) or the trained leg (P > 0.999). MG sarcomere length also showed an effect of training (P = 0.008), with no group (P = 0.267) or interaction effects (P = 0.594). The LG fascicle length showed a training × group interaction (P = 0.002) where, compared to the untrained leg, fascicle length in the trained leg was longer in Burst (+12.5 ± 5.9%, ES = 1.89; P < 0.001), but not in Iso (+3.2 ± 4.9%, ES = 0.57; P = 0.416). There were no differences in fascicle length between groups in either the untrained (P = 0.792) or the trained leg (P = 0.565). Sarcomere length in the LG did not exhibit any significant effects of training (0.782), group (P = 0.097) or any interaction (P = 0.217). It is important to note that, unlike SSN, the measured fascicle and sarcomere lengths will depend on the length at which the muscle was fixed at. Therefore, any imprecision in the joint angles the legs were fixed at could impact these values, and complicate any interpretations made from these data.

**Supplemental Figure 2.**
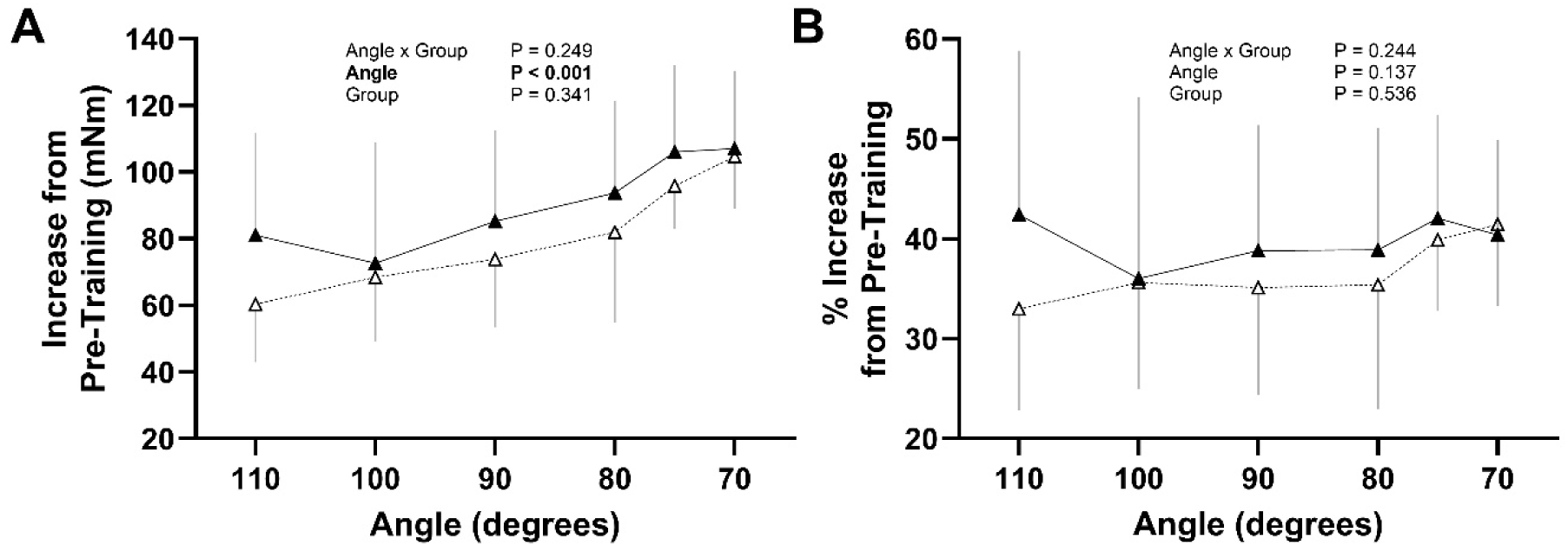
Absolute (Panel A) and Percentage (Panel B) improvements in active torque across the ankle angles tested. **Results:** As evident by the significant angle × training interaction in the 3-Way ANOVA (see Figure 2 in main file), the absolute improvement in active torque from pre-training increased with muscle length (Panel A). However, when expressed as a percentage increase (Panel B), this length-dependent effect was not apparent.

## REFERENCES

1. Jorgenson KW, Phillips SM, Hornberger TA. Identifying the structural adaptations that drive the mechanical load-induced growth of skeletal muscle: a scoping review. Cells. 2020;9(7):1658.

2. Hinks A, Hawke TJ, Franchi MV, Power GA. The importance of serial sarcomere addition for muscle function and the impact of aging. J Appl Physiol. 2023;135(2):375–393. doi:10.1152/japplphysiol.00205.2023

3. Butterfield TA, Herzog W. The magnitude of muscle strain does not influence serial sarcomere number adaptations following eccentric exercise. Pflüg Arch – Eur J Physiol. 2006;451(5):688–700. doi:10.1007/s00424-005-1503-6

4. Goldspink G. Malleability of the motor system: a comparative approach. J Exp Biol. 1985;115(1):375–391. doi:10.1242/jeb.115.1.375

5. De Lima-E-Silva FX, Oliveira GS, Medeiros TM, Dornelles MP, Ribeiro-Alvares JBA, Baroni BM. Flexibility, strength, and fascicle length of football players with and without history of hamstring strain injury in the prior season. Sci Med Footb. 2020;4(4):322–328. doi:10.1080/24733938.2020.1751871

6. Andrews MH, Shield AJ, Lichtwark GA, Pincheira PA. Hamstring Injury Mechanisms and Eccentric Training-Induced Muscle Adaptations: Current Insights and Future Directions. Sports Med. 2025;55(10):2429–2443. doi:10.1007/s40279-025-02291-6

7. Abe T, Kumagai K, Brechue WF. Fascicle length of leg muscles is greater in sprinters than distance runners. Med Sci Sports Exerc. 2000;32(6):1125–1129.

8. Cooper AN, McDermott WJ, Martin JC, Dulaney SO, Carrier DR. Great power comes at a high (locomotor) cost: the role of muscle fascicle length in the power versus economy performance trade-off. J Exp Biol. 2021;224(21):jeb236679. doi:10.1242/jeb.236679

9. Kumagai K, Abe T, Brechue WF, Ryushi T, Takano S, Mizuno M. Sprint performance is related to muscle fascicle length in male 100-m sprinters. J Appl Physiol. 2000;88(3):811–816. doi:10.1152/jappl.2000.88.3.811

10. Lee HJ, Lee KW, Takeshi K, Lee YW, Kim HJ. Correlation analysis between lower limb muscle architectures and cycling power via ultrasonography. Sci Rep. 2021;11(1):5362.

11. Blazevich AJ, Herzog W, Nunes JP. Triggering sarcomerogenesis: Examining key stimuli and the role attributed to eccentric training—Historical, systematic, and meta-analytic review. J Sport Health Sci. Published online 2025:101073.

12. Uçar N, Öner H, Kuş MA, Karaca H, Fırat T. The effect of neuromuscular electrical stimulation applied at different muscle lengths on muscle architecture and sarcomere morphology in rats. Anat Rec. 2024;307(2):356–371. doi:10.1002/ar.25240

13. Rilling A, Zero AM, Kirkup AQ, et al. Contraction type dependence of serial sarcomere addition following resistance training. BioRxiv. Preprint posted online 2026. doi:10.64898/2026.09.29.755478

14. Blazevich AJ, Herzog W, Nunes JP. Sarcomerogenesis: Evidence, context and key Stimuli–Response to the commentary by Power et al. J Sport Health Sci. Published online 2025:101090.

15. Power GA, Franchi MV, Hinks A. The mechanical loading environment associated with eccentric exercise is one of the key stimuli to trigger sarcomerogenesis: It’sa stretch to say eccentric exercise does not promote serial sarcomerogenesis. J Sport Health Sci. Published online 2025:101089.

16. Butterfield TA, Herzog W. Quantification of muscle fiber strain during in vivo repetitive stretch-shortening cycles. J Appl Physiol. 2005;99(2):593–602. doi:10.1152/japplphysiol.01128.2004

17. Sharifnezhad A, Marzilger R, Arampatzis A. Effects of load magnitude, muscle length and velocity during eccentric chronic loading on the longitudinal growth of the vastus lateralis muscle. J Exp Biol. 2014;217(15):2726–2733.

18. Hinks A, Rilling A, Rand C, Borkowski M, Power GA. Construction of Constant-Load (Isotonic) and Constant-Velocity (Isokinetic) Torque-Velocity-Power Profiles In vivo for the Rat Plantar Flexors. J Vis Exp JoVE. 2025;(224):e68858.

19. Ashida Y, Himori K, Tatebayashi D, Yamada R, Ogasawara R, Yamada T. Effects of contraction mode and stimulation frequency on electrical stimulation-induced skeletal muscle hypertrophy. J Appl Physiol. 2018;124(2):341–348. doi:10.1152/japplphysiol.00708.2017

20. MacIntosh BR, MacNaughton MB. The length dependence of muscle active force: considerations for parallel elastic properties. J Appl Physiol. 2005;98(5):1666–1673. doi:10.1152/japplphysiol.01045.2004

21. Hill AV. The heat of shortening and the dynamic constants of muscle. Proc R Soc Lond Ser B – Biol Sci. 1938;126(843):136–195. doi:10.1098/rspb.1938.0050

22. Ward SR, Lieber RL. Density and hydration of fresh and fixed human skeletal muscle. J Biomech. 2005;38(11):2317–2320.

23. Zuurbier CJ, Heslinga JW, Lee-de Groot MBE, Van der Laarse WJ. Mean sarcomere length-force relationship of rat muscle fibre bundles. J Biomech. 1995;28(1):83–87. doi:10.1016/0021-9290(95)80009-3

24. Hedges LV. Effect sizes for experimental research. Br J Math Stat Psychol. 2026;79(1):31–45. doi:10.1111/bmsp.12389

25. Cohen J. Statistical Power Analysis for the Behavioural Sciences. Hillside. 1988.

26. Rilling A, Kirkup AQ, Hinks A, Franchi MV, Power GA. Serial sarcomere adaptation to eccentric overload training is blunted in ovariectomized rats. Physiol Rep. 2026;14(16):e71070. doi:10.14814/phy2.71070

27. Rilling A, Hinks A, Kirkup AQ, Franchi MV, Power GA. Eccentric overload training promotes serial sarcomerogenesis to a greater extent than conventional resistance training. Am J Physiol-Cell Physiol. 2026;330(2):C345–C355. doi:10.1152/ajpcell.00713.2025

28. Hinks A, Patterson MA, Njai BS, Power GA. Age-related blunting of serial sarcomerogenesis and mechanical adaptations following 4 wk of maximal eccentric resistance training. J Appl Physiol. 2024;136(5):1209–1225. doi:10.1152/japplphysiol.00041.2024

29. Hinks A, Vlemmix E, Power GA. Submaximal eccentric resistance training increases serial sarcomere number and improves dynamic muscle performance in old rats. Physiol Rep. 2024;12(19):e70036. doi:10.14814/phy2.70036

30. Hutchison DL, Roy RR, Hodgson JA, Edgerton VR. EMG amplitude relationships between the rat soleus and medial gastrocnemius during various motor tasks. Brain Res. 1989;502(2):233–244.

31. Chen J, Mashouri P, Fontyn S, et al. The influence of training-induced sarcomerogenesis on the history dependence of force. J Exp Biol. Published online January 1, 2020:jeb.218776. doi:10.1242/jeb.218776

32. Pelland JC, Remmert JF, Robinson ZP, Hinson SR, Zourdos MC. The Resistance Training Dose Response: Meta-Regressions Exploring the Effects of Weekly Volume and Frequency on Muscle Hypertrophy and Strength Gains. Sports Med. 2026;56(2):481–505. doi:10.1007/s40279-025-02344-w

33. Rader EP, Naimo MA, Layner KN, et al. Enhancement of Skeletal Muscle in Aged Rats Following High-Intensity Stretch-Shortening Contraction Training. Rejuvenation Res. 2017;20(2):93–102. doi:10.1089/rej.2016.1816

34. Ogasawara R, Arihara Y, Takegaki J, Nakazato K, Ishii N. Relationship between exercise volume and muscle protein synthesis in a rat model of resistance exercise. J Appl Physiol. 2017;123(4):710–716. doi:10.1152/japplphysiol.01009.2016

35. Kobayashi K, Ogasawara R, Tsutaki A, Lee K, Ochi E, Nakazato K. Genetic strain-dependent protein metabolism and muscle hypertrophy under chronic isometric training in rat gastrocnemius muscle. Physiol Res. 2012;61(5):527.

36. Cutlip RG, Baker BA, Geronilla KB, et al. Chronic exposure to stretch–shortening contractions results in skeletal muscle adaptation in young rats and maladaptation in old rats. Appl Physiol Nutr Metab. 2006;31(5):573–587. doi:10.1139/h06-033

37. Adams GR, Hather BM, Baldwin KM, Dudley GA. Skeletal muscle myosin heavy chain composition and resistance training. J Appl Physiol. 1993;74(2):911–915. doi:10.1152/jappl.1993.74.2.911

38. Caiozzo VJ, Haddad F, Baker MJ, Baldwin KM. Influence of mechanical loading on myosin heavy-chain protein and mRNA isoform expression. J Appl Physiol. 1996;80(5):1503–1512. doi:10.1152/jappl.1996.80.5.1503

39. Bottinelli R, Betto R, Schiaffino S, Reggiani C. Unloaded shortening velocity and myosin heavy chain and alkali light chain isoform composition in rat skeletal muscle fibres. J Physiol. 1994;478(2):341–349. doi:10.1113/jphysiol.1994.sp020254

40. Ranatunga KW, Thomas PE. Correlation between shortening velocity, force-velocity relation and histochemical fibre-type composition in rat muscles. J Muscle Res Cell Motil. 1990;11(3):240–250. doi:10.1007/BF01843577

41. MacIntosh BR, Herzog W, Suter E, Wiley JP, Sokolosky J. Human skeletal muscle fibre types and force: velocity properties. Eur J Appl Physiol. 1993;67(6):499–506. doi:10.1007/BF00241645

42. Wackerhage H, Schoenfeld BJ, Hamilton DL, Lehti M, Hulmi JJ. Stimuli and sensors that initiate skeletal muscle hypertrophy following resistance exercise. J Appl Physiol. 2019;126(1):30–43. doi:10.1152/japplphysiol.00685.2018

43. Egan B, Sharples AP. Molecular responses to acute exercise and their relevance for adaptations in skeletal muscle to exercise training. Physiol Rev. 2023;103(3):2057–2170. doi:10.1152/physrev.00054.2021

44. Roberts MD, McCarthy JJ, Hornberger TA, et al. Mechanisms of mechanical overload-induced skeletal muscle hypertrophy: current understanding and future directions. Physiol Rev. 2023;103(4):2679–2757. doi:10.1152/physrev.00039.2022

45. Franchi MV, Reeves ND, Narici MV. Skeletal muscle remodeling in response to eccentric vs. concentric loading: morphological, molecular, and metabolic adaptations. Front Physiol. 2017;8:447.

46. Proske U, Morgan DL. Muscle damage from eccentric exercise: mechanism, mechanical signs, adaptation and clinical applications. J Physiol. 2001;537(2):333–345. doi:10.1111/j.1469-7793.2001.00333.x

47. Davis JF, Khir AW, Barber L, et al. The mechanisms of adaptation for muscle fascicle length changes with exercise: Implications for spastic muscle. Med Hypotheses. 2020;144:110199.

48. Hibbert JE, Jorgenson KW, Sayed RKA, et al. Mechanical loading induces the longitudinal growth of muscle fibers via a rapamycin-insensitive mechanism. Sci Adv. 2026;12(7):eaec5134. doi:10.1126/sciadv.aec5134

49. Franchi MV, Ruoss S, Valdivieso P, et al. Regional regulation of focal adhesion kinase after concentric and eccentric loading is related to remodelling of human skeletal muscle. Acta Physiol. 2018;223(3):e13056. doi:10.1111/apha.13056

50. Ingber DE. From Cellular Mechanotransduction to Biologically Inspired Engineering: 2009 Pritzker Award Lecture, BMES Annual Meeting October 10, 2009. *Ann Biomed Eng*. 2010;38(3):1148–1161. doi:10.1007/s10439-010-9946-0

51. Taylor DC, Dalton JD, Seaber AV, Garrett WE. Viscoelastic properties of muscle-tendon units: The biomechanical effects of stretching. Am J Sports Med. 1990;18(3):300–309. doi:10.1177/036354659001800314

52. Burkholder TJ. Permeability of C2C12 myotube membranes is influenced by stretch velocity. Biochem Biophys Res Commun. 2003;305(2):266–270.

53. Van Der Pijl R, Strom J, Conijn S, et al. Titin based mechanosensing modulates muscle hypertrophy. J Cachexia Sarcopenia Muscle. 2018;9(5):947–961. doi:10.1002/jcsm.12319

54. Brynnel A, Hernandez Y, Kiss B, et al. Downsizing the molecular spring of the giant protein titin reveals that skeletal muscle titin determines passive stiffness and drives longitudinal hypertrophy. elife. 2018;7:e40532.

55. Granzier H, Labeit D, Wu Y, Labeit S. Titin as a modular spring: emerging mechanisms for elasticity control by titin in cardiac physiology and pathophysiology. In: Linke WA, Granzier H, Kellermayer MSZ, eds. Mechanics of Elastic Biomolecules. Springer Netherlands; 2003:457–471. doi:10.1007/978-94-010-0147-2_7

56. Bloch RJ, Gonzalez-Serratos H. Lateral force transmission across costameres in skeletal muscle. Exerc Sport Sci Rev. 2003;31(2):73–78.

57. Fortuna R, Power GA, Mende E, Seiberl W, Herzog W. Residual force enhancement following shortening is speed-dependent. Sci Rep. 2016;6(1):21513.

58. Herzog W, Leonard TR. The history dependence of force production in mammalian skeletal muscle following stretch-shortening and shortening-stretch cycles. J Biomech. 2000;33(5):531–542. doi:10.1016/S0021-9290(99)00221-3

